# Systematic Evaluation of Feature Representations for Cancer-Associated sORF Prediction in Non-coding RNA

**DOI:** 10.64898/2026.06.16.732659

**Authors:** Fabiana Rodrigues de Goes, Makanaka Mazheke, Amanda Piveta Schnepper, Aparajita Karmakar, Nayane de Souza, Robson Francisco Carvalho, Mark Basham, Alexandre Rossi Paschoal

## Abstract

Short open reading frames (sORFs) within non-coding RNAs (ncRNAs) have arisen as a hidden layer of gene regulation, encoding small peptides that represent a new class of cancer regulators with diagnostic and therapeutic potential. However, inferring associations between sORFs to specific cancer types remains challenging and requires computational approaches for accurate prediction. Recently, the CoraL framework introduced the first computational approach for predicting cancer-associated peptides, focusing primarily on model architecture while overlooking how feature extraction strategies influence predictive accuracy. We present a systematic evaluation of machine learning models and feature extraction approaches to predict cancer-associated sORFs across 15 cancer types. We benchmarked seven traditional machine learning algorithms combined with three feature extraction methods: k-mer frequency, Word2Vec embeddings, and genomic language model (gLM)-based embeddings. To our knowledge, this is the first study applying gLM-derived embeddings to the prediction of cancer-associated sORFs in ncRNA. Our results show that traditional machine learning models with appropriate feature extraction outperform the CoraL baseline across all cancer types, achieving up to 10% higher accuracy in some of the 15 evaluated datasets. Interestingly, k-mer features consistently outperformed gLM embeddings without fine-tuning, suggesting that local sequence composition may provide more discriminative information for this task and that pre-trained genomic representations may require task-specific adaptation to fully capture these patterns. Additionally, we observed that the way sequences are tokenized, such as the k-mer length, can affect performance: longer fragments (e.g., k=7) sometimes reduced accuracy for Random Forest but had a smaller effect on MLP. Our findings suggest that appropriate feature engineering can provide greater improvements than increasing model complexity.

## 1 Introduction

Non-coding RNAs (ncRNAs) have traditionally been defined as RNA molecules that are transcribed with low or no protein-coding capacity [1–4]. However, not surprisingly, the emergence of ribosome profiling (Ribo-Seq) technique in 2009 [5] opened a hidden layer of what constitutes coding or non-coding capacity by revealing that non-canonical genomic regions—including untranslated regions, introns, and ncRNAs—contain small open reading frames (sORFs) with the ability to be translated into functional small peptides (sPEPs) [6, 7]. This discovery has broken the paradigm beyond traditional proteins, prompting recent efforts by international consortia to establish standardized ORF annotation [8] and definitions for translated regions, termed ”translons” [9, 10]. Evidence now demonstrates that ncRNA-encoded small peptides (ncPEPs) are not translational artifacts [5, 11–13], but rather *bona fide* functional molecules involved in diverse biological processes, including disease pathogenesis and cancer [1, 6, 7, 14, 15]. For example, the dysregulated expression of specific ncPEPs contributes to cancer initiation and progression, highlighting their potential as biomarkers and therapeutic targets [14, 15].

Ribo-Seq, mass spectrometry, and other protein detection strategies currently represent the gold standard experimental platform for discovering and validating sPEPs [6, 16]. However, these approaches are resource-intensive and require substantial time, costs, and specialized laboratory infrastructure [6, 8, 16]. Moreover, both methods face inherent technical and biological limitations, including detection sensitivity thresholds, tissue-specific expression patterns, and transient peptide stability, restricting their capacity to capture the full spectrum of functional sPEPs within cells [17]. Consequently, a substantial fraction of potentially functional small peptides remains undetected, creating a critical knowledge gap in our understanding of this regulatory layer. In response to these limitations, and in the current era of artificial intelligence, there is an urgent need for open-access computational tools that can systematically predict and characterize cancer-relevant sORFs, particularly within ncRNAs. By complementing experimental approaches and prioritizing candidate sORFs for downstream validation, AI-based methods have the potential to accelerate the identification of functional sPEPs and advance our understanding of their roles in cancer biology.

In this context, machine learning approaches have emerged as powerful tools for predicting sORFs and inferring the sPEPs they encode. However, most previous studies have focused mainly on detecting sORFs, without exploring their potential biological or clinical relevance [18–20]. To overcome this limitation, CoraL [21] was the first framework designed to predict associations between cancer and both ncPEPs and sORFs, based on a contrastive learning, TextCNN [22], and meta-learning approach. However, for cancer-associated sORF prediction, the framework relies on a TextCNN-based sequence classifier, leaving unexplored the potential impact of alternative, tokenization, feature representations and learning algorithms. Moreover, CoraL does not systematically evaluate how different sequence encoding strategies influence predictive performance.

Unlike natural language, DNA sequences lack inherent structure, spacing, or semantic boundaries, making tokenization a crucial step for transforming raw sequences into meaningful computational inputs. A common approach is the use of k-mers, which segment sequences into fragments of length *k* to capture local compositional patterns while preserving biologically relevant information. Beyond k-mer encoding, other representation learning methods can be applied to learn embedding vectors, and the choice of tokenization or feature extraction strategy can significantly influence model performance [23]. Moreover, because no single algorithm or feature representation consistently achieves superior performance across all datasets and prediction tasks, a systematic evaluation of different feature representations and learning algorithms is crucial to identify the optimal combination for a given problem.

In this study, we propose an analysis to address the challenge of predicting cancer-associated sORFs within ncRNA across 15 cancer types. To this end, we evaluated the performance of seven well-established classifiers: Random Forest (RF) [24], Support Vector Machine (SVM) [25], Multilayer Perceptron (MLP) [26], Logistic Regression (LR), XGBoost, Naive Bayes (NB), and k-nearest neighbors (KNN). We also compared three distinct feature extraction strategies: k-mer frequency, Word2Vec embeddings, and genomic language model (gLM)-based embeddings. To our knowledge, this is the first study applying an LLM-based approach to this specific application. Our findings indicate clear differences in how these representations influence classification performance, with simpler feature extraction approaches, such as k-mer, frequently achieving more stable and accurate results than gLM-derived embeddings. Furthermore, when benchmarked against CoraL our most accurate models achieved higher accuracy across all datasets. In summary, this work offers a comprehensive end-to-end evaluation of feature representation strategies for cancer-associated sORF within ncRNA prediction and demonstrates that carefully designed feature representations can enable conventional machine learning models to outperform more complex modeling approaches.

## 2 Materials and Methods

The workflow consisted of two-step data analysis: (I) feature extraction, and (II) classification phases (Figure 1). The sequence data were transformed into numerical representations using three methods: k-mer frequency, Word2Vec embeddings, and pre-trained gLM embeddings. These features were used to train Random Forest, Support Vector Machine, Multilayer Perceptron, Logistic Regression, XGBoost, Naive Bayes and KNN classifiers with hyperparameters optimized via 3-fold cross-validation. Model performance on held-out test data was evaluated using standard classification metrics (accuracy, specificity, and sensitivity).

**Figure 1:**
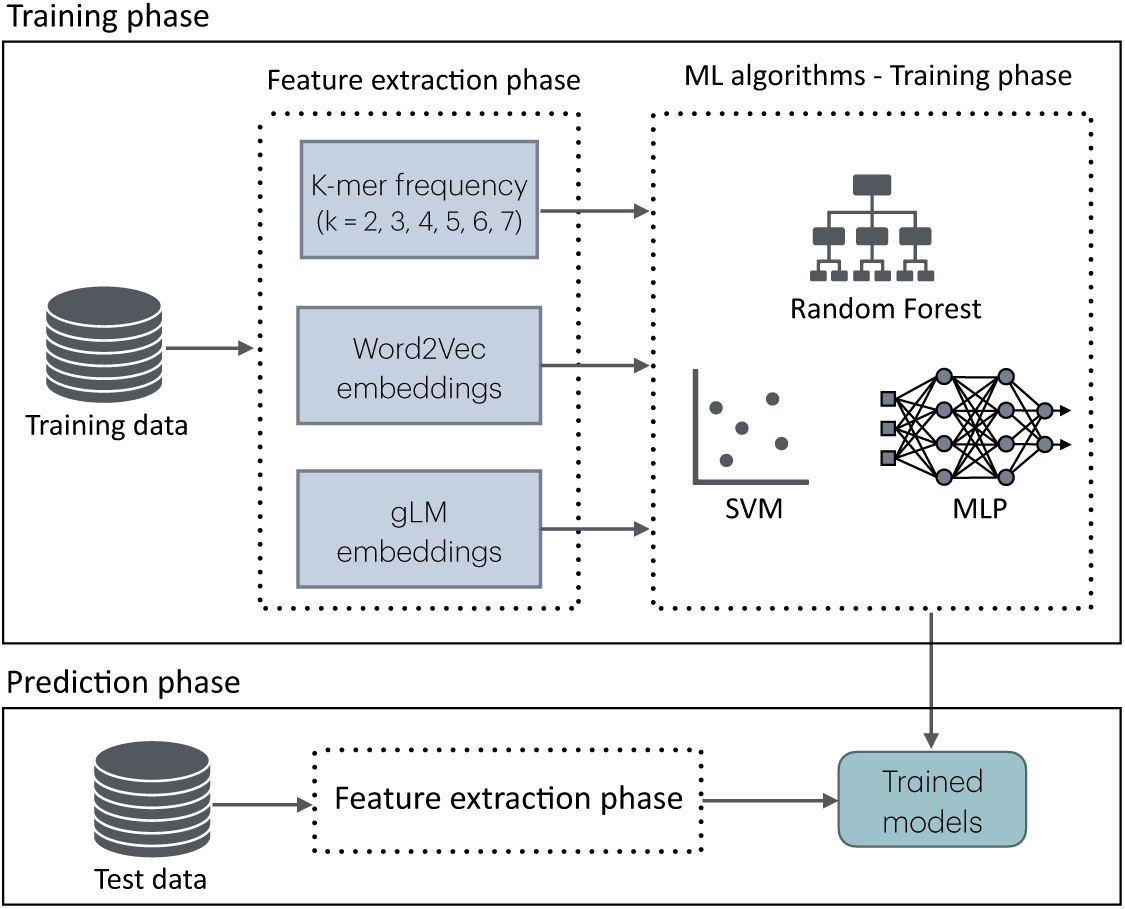
The overall workflow is based on a training phase that involves three feature strategies with three ML algorithms and the predictive phase, in which the trained model is used to predict if a given sORF is associated with a cancer in ncRNA data.

### 2.1 Datasets

SPENCER [27] is a comprehensive repository of experimentally verified ncPEPs and their corresponding sORFs across multiple cancer types. Using SPENCER as a source, the CoraL authors extracted cancer-associated sORFs, covering 15 distinct cancer types, and established standardized training and test splits suitable for computational modeling (Figure S1). We utilized the 15 cancer-specific sORF datasets created by the CoraL framework [21], which were derived from the SPENCER database [27].

Because SPENCER contains only experimentally validated peptides and their sORFs (positive samples), the CoraL study generated synthetic negative samples by shuffling the nucleotide sequences of authentic sORFs. This procedure produced balanced datasets with the same number of positive and negative instances. The original publications provide comprehensive details on the curation methodologies of SPENCER [27] and CoraL [21]. In our work, we directly adopted the 15 datasets provided by CoraL [21], including their original training and test partitions, to ensure comparability with the performance reported in their study.

### 2.2 Feature extraction approaches

We applied three encoding strategies to convert them into numerical representations: (i) k-mer frequency encoding, (ii) Word2Vec embeddings, and (iii) gLMs. For each sequence, we calculated k-mer frequencies using *k* values ranging from 2 to 7, where the total number of possible overlapping *k*-mers was calculated as

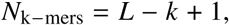

where *L* is the sequence length and *k* is the length of each *k*-mer. To mitigate the effects of varying sequence lengths, raw k-mer counts were converted into relative frequencies. Specifically, for each sequence *i* and each k-mer *j* , the count *c*_*i*,_ *j* was normalized by the total number of k-mers in the sequence, resulting in a relative frequency *f*_*i*, *j*_ :

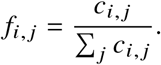

Word2Vec Skip-gram model [28] was employed to map k-mers to a numerical space. For each dataset, a Word2Vec model was trained using all sequences from the training set. The sequences were first tokenized into k-mers to capture local sequence context. Next, the Word2Vec model was used to generate embeddings for all sequences, including both training and test sets. The final embedding vector for each sequence was obtained by averaging the embedding vectors of all its constituent k-mers. The resulting embeddings can be employed as input features for training a classification model or used as a classification model.

For the gLM-based approach, we employed two pretrained models: DNABERT2 [29] and the Nucleotide Transformer (NT) [30]. Both models are transformer-based architectures trained on large-scale genomic data to capture contextual dependencies within DNA sequences. Each input sequence was tokenized according to the model’s vocabulary and processed through the model to obtain hidden representations. The classify token ([CLS]), used in transformer architectures to summarize the input sequence, was considered to obtain a fixed-length representation. The embedding associated with this token was extracted from the last hidden layer to represent the global contextual information of the sequence. Additionally, alternative methods for processing gLM embeddings, such as average and maximum pooling, were adopted to aggregate contextual information across all tokens in the sequence.

### 2.3 Experimental Analysis

In this study, training and test sets were used in distinct phases to ensure unbiased evaluation of model performance. The training set was used for hyperparameter optimization of the seven machine learning algorithms, with the search space defined by the hyperparameter grids (Table 1). The model selection was performed using 3-fold cross-validation. The same hyperparameter grids were consistently applied across all three sequence representation strategies evaluated in this work.

**Table 1:** Hyperparameter settings for each machine learning algorithm used in this work.

| Algorithm | Hyperparameters | Regularization | Implementation |
| --- | --- | --- | --- |
| Random Forest | Trees: 200, 300, 400 | – | scikit-learn<br>RandomForestClassifier |
| SVM | Kernel: linear, rbf | $C$: 0.1, 1, 10 | scikit-learn SVC |
| MLP | Hidden layers: (32,16), (100) | L2 penalty ($\alpha$): $10^{-4}$, $10^{-3}$ | scikit-learn MLPClassifier |
| Logistic Regression | Inverse regularization strength ($C$): 0.01, 0.1, 1, 10 | L1, L2 penalty | scikit-learn<br>LogisticRegression |
| XGBoost | Trees: 200, 300, 400 | Learning rate ($\eta$): 0.01, 0.1 | XGBoost XGBClassifier |
| KNN | $K$: 3, 5, 7, 10 | – | scikit-learn<br>KNeighborsClassifier |
| Naive Bayes | Variance smoothing: $10^{-9}$, $10^{-8}$, $10^{-7}$ | – | scikit-learn GaussianNB |

*Note:* All hyperparameters not explicitly specified in this table were set to their default values as defined in the respective libraries.
All models were implemented using scikit-learn (v1.7.2) and XGBoost (v3.1.3).

Additionally, we optimized the tokenization parameters for k-mer–based representations to determine the optimal *k* value. This parameter was tuned for both the k-mer frequency representation and Word2Vec-based embeddings. For the Word2Vec approach, the embedding vector size ([150, 250, 300]) was also tuned to assess its influence on downstream classification performance. Regarding gLM–based embeddings, we extracted generic representations from the pretrained DNABERT2 and NT models, without any task-specific fine-tuning. These embeddings were used as general-purpose representations of genomic sequences and served as input features for the ML algorithms to evaluate the generalization capacity of pretrained models across cancer types.

Within the training set, a stratified 3-fold cross-validation (CV) procedure was applied to both hyperparameter analysis and model selection. First, CV was used to analyse the influence of hyperparameter variations on model performance. Specifically, we computed the average results across all 15 cancer-specific datasets for each combination of algorithms and hyperparameters, considering different values of *k* and embedding vector size (just for the Word2Vec-based approach). This comprehensive analysis enabled the identification of the hyperparameter configuration that yielded the most consistent overall performance across datasets. Once the optimal hyperparameter configuration was established, it was fixed for subsequent experiments to evaluating the specific effects of representation parameters (*k* and embedding vector size) on model accuracy. Finally, the best hyperparameter combination was identified for each of the 15 cancer datasets and used to retrain the corresponding classification model on the full training set. Each model was evaluated on its respective held-out test set, and the results were compared against the CoraL baseline. This protocol ensured that the test data remained completely unseen throughout model development, providing a robust and unbiased estimate of generalization performance.

## 3 Results

### 3.1 Analysis of Hyperparameter Effects

To assess the influence of hyperparameters on model performance, we generated box plots showing the average results for each combination of algorithm and hyperparameter in all 15 cancer-specific datasets and for each feature type: k-mer (Figure 2a), Word2Vec (Figure 2b), DNABERT2 (Figure 3a) and NT (Figure 3b). Some algorithms exhibited consistent behavior across all sequence representations, indicating low sensitivity to hyperparameter selection. Random Forest was the most stable classifier overall, the ensemble size had limited influence on predictive performance within the evaluated range and the highest accuracies were acheived with k-mer and Word2Vec features. A similar pattern was observed for Naive Bayes, whose performance remained nearly unchanged across the evaluated smoothing hyperparameter for all feature representations. MLP also demonstrated low hyperparameter sensitivity, while some variation was observed across hidden-layer configurations and learning rates, the median accuracies remained stable for all feature extraction methods, particularly for Word2Vec and transformer-based embeddings. Unlike Random Forest and MLP, XGBoost exhibited a consistent increase in performance with increasing numbers of estimators and learning rates across all feature representations.

**Figure 2:**
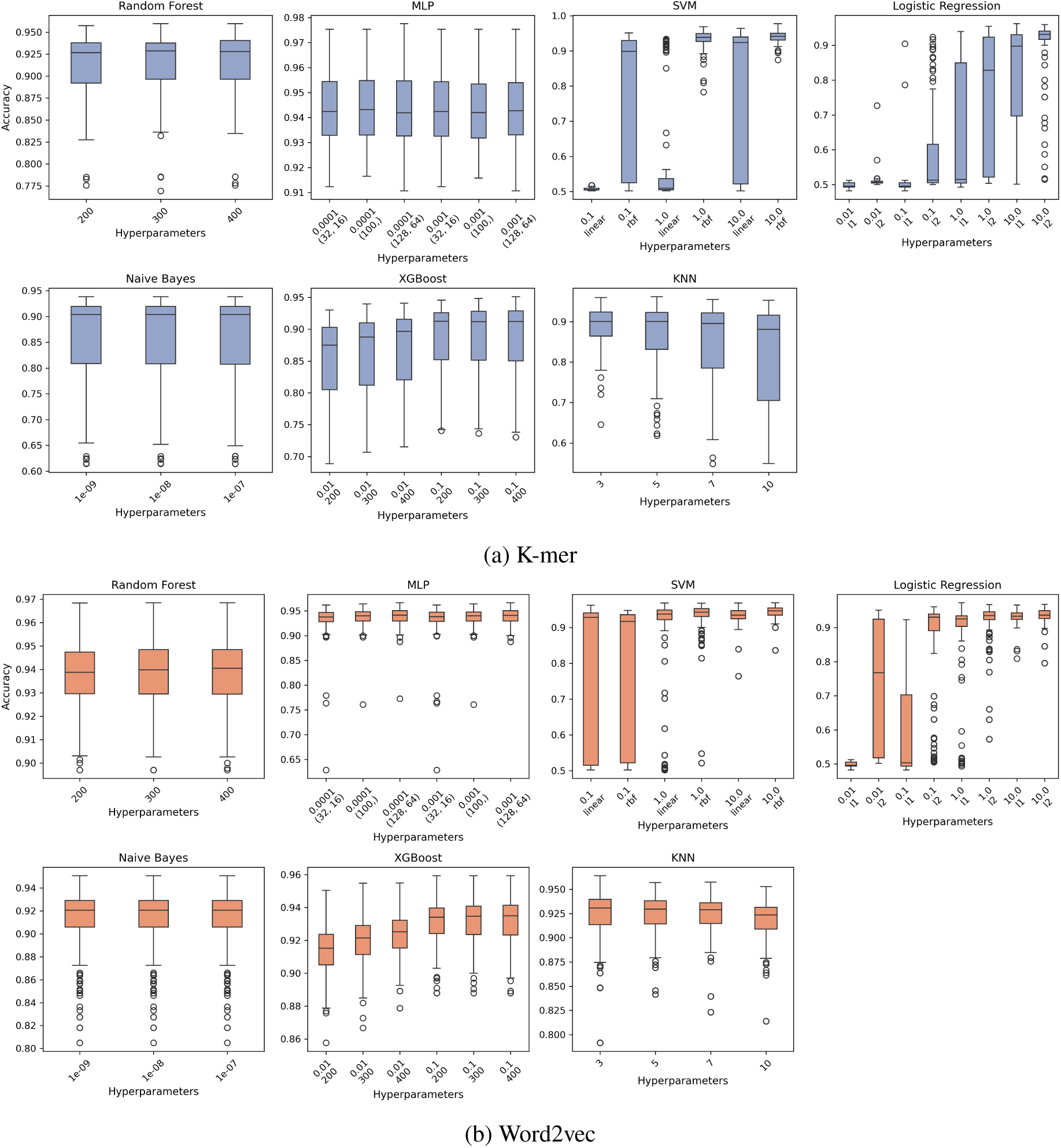
Hyperparameter performance analysis across feature representation strategies. Boxplots show the mean cross-validation accuracies for each classifier using (a) k-mer and (b) Word2Vec representations under varying hyperparameter settings.

**Figure 3:**
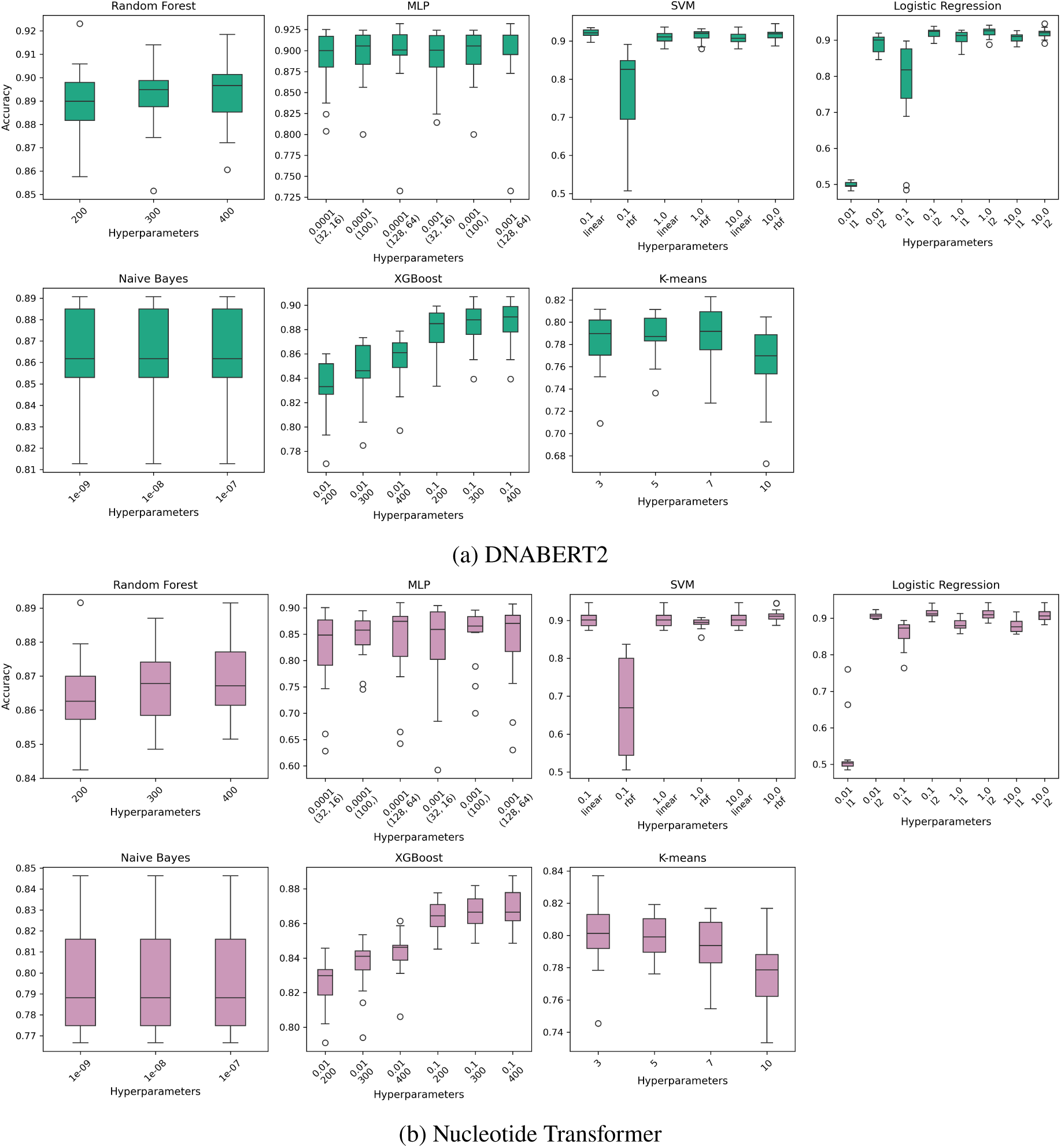
Hyperparameter performance analysis across feature representation strategies. Boxplots showing average cross-validation accuracies for each classifier using (a) DNABERT2 and (b) NT under different hyperparameter settings.

In contrast, SVM and Logistic Regression exhibited the strongest dependence on hyperparameter selection. However, the impact of hyperparameter selection varied according to the feature representation. For k-mer and Word2Vec features, SVM and Logistic Regression exhibited considerable sensitivity to kernel choice and regularization strength, respectively. In comparison, for DNABERT2 and NT embeddings, the performance distributions became considerably more homogeneous among the hyperparameter configurations. This suggests that transformer-derived embeddings provided a feature space in which these two algorithms were less sensitive to hyperparameter tuning.

The behavior of KNN also depended on the feature representation. The smallest performance variation across neighborhood sizes was observed for Word2Vec embeddings, where all values of *k* achieved similar median accuracies. In contrast, k-mer and transformer-based embeddings showed greater sensitivity to the choice of neighborhood size, particularly for larger values of *k*.

To investigate the effects of representation parameters on model accuracy, we analyzed the variation in performance across different *k* values for each combination of feature representation and classification algorithm on the 15 cancer datasets (Figure 4). The combination of RF with k-mer features exhibited the largest variability in performance, with a consistent decrease in accuracy for *k* values ≥ 5. In contrast, Word2Vec-based representations were generally more stable across *k* variations. The five smallest datasets in terms of sample size, Thyroid, Tongue, Ovary, Liver, and Gastric (listed from smallest to largest) exhibited the most pronounced fluctuations in their performance curves. In comparison, larger datasets displayed more tightly grouped curves, indicating reduced sensitivity to the choice of *k*. These observations suggest that smaller datasets are more susceptible to variations in representation parameters, particularly when using k-mer features with RF.

**Figure 4:**
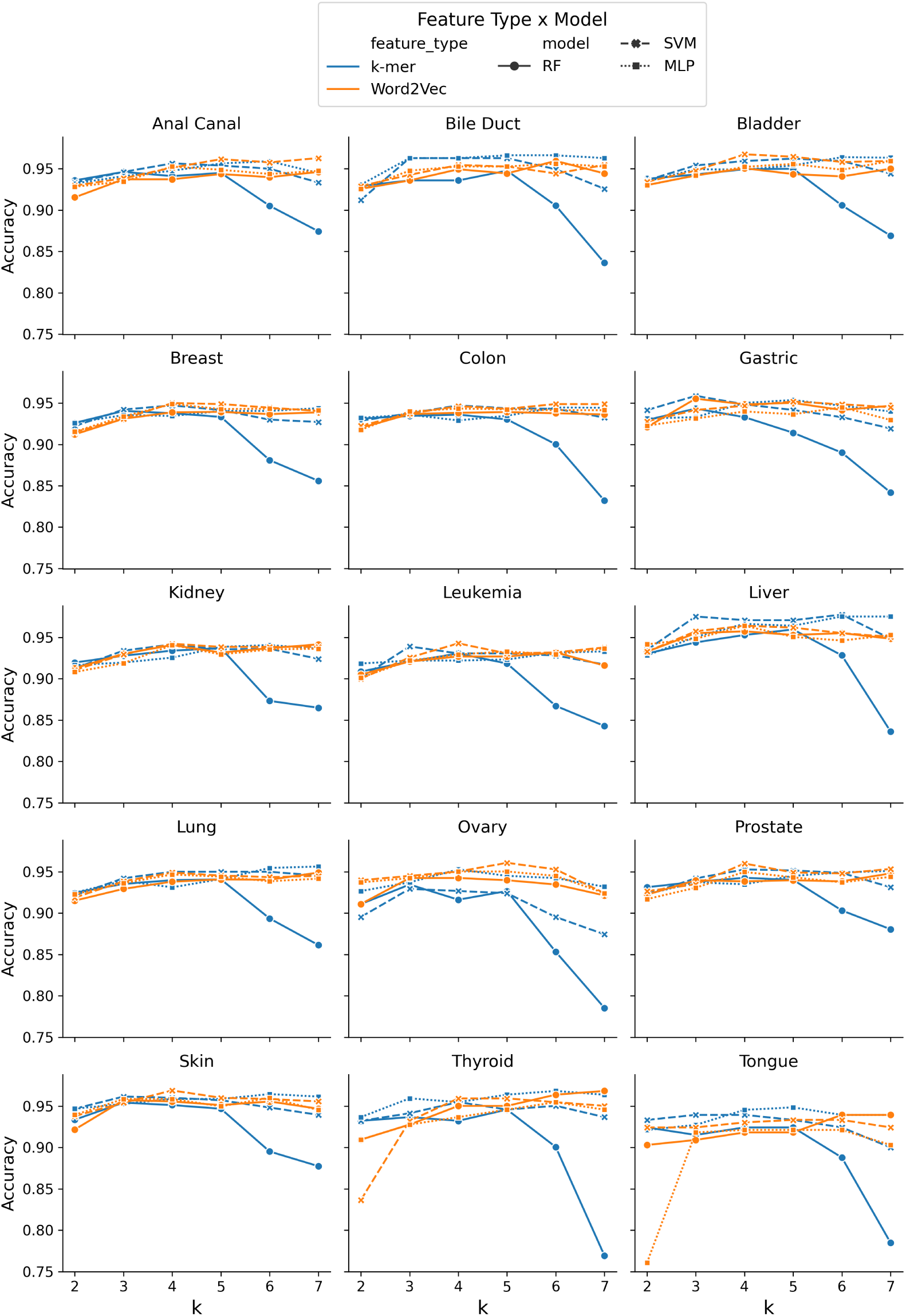
Performance variation across *k* values for different feature representations and classification algorithms in the 15 cancer datasets. Each line represents a specific combination of feature type and algorithm.

### 3.2 Analysis of Pooling Effects

The comparison of pooling strategies demonstrated that the choice of embedding aggregation method affected the downstream classification performance for both DNABERT2 (Figure 5a) and NT (Figure 5b). Overall, mean pooling consistently achieved the best or among the best accuracy values across most cancer types and the seven models. This behavior suggests that averaging token-level contextual representations provides a more stable and informative characterization of the sequence than CLS or max pooling.

**Figure 5:**
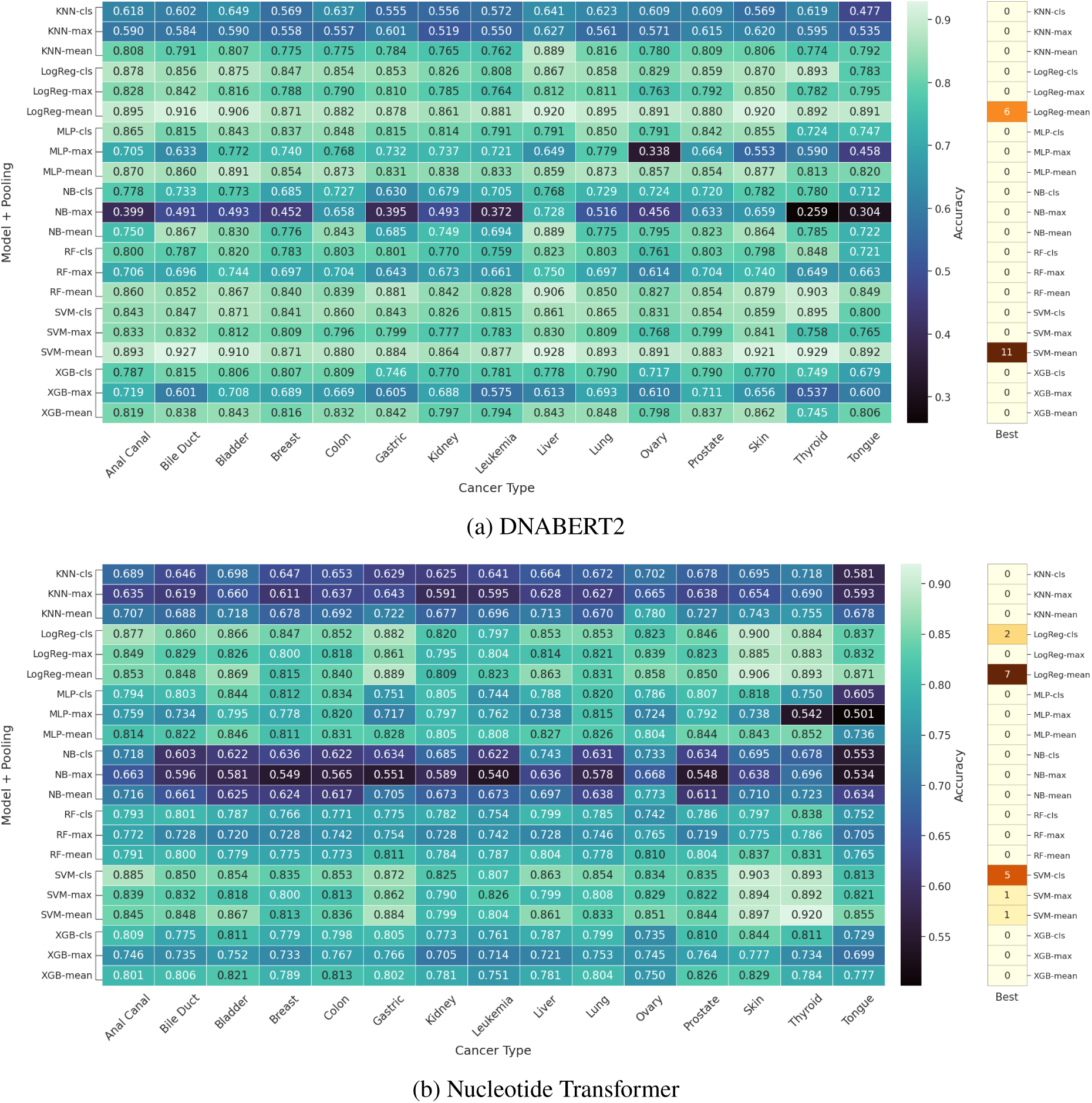
Heatmaps showing the highest cross-validation accuracies obtained by different machine learning models combined with distinct embedding pooling strategies (CLS, max, and mean) across the datasets. The ’Best’ column reports the number of times each model–pooling combination achieved the highest accuracy across the 15 datasets (ties counted for all winners). Heatmaps (a) and (b) correspond to embeddings generated by DNABERT2 and NT, respectively.

The effect of pooling strategy was evident for DNABERT2, where the highest accuracies were predominantly obtained using mean-pooled embeddings (Figure 5a and Figure S5a). Although mean pooling also achieved the best overall results for NT, CLS pooling produced comparable accuracies particularly when combined with SVM and Logistic Regression (Figure 5b and S5b). The reduced performance gap between these aggregation strategies suggests that the CLS representation derived from NT may retain global sequence information more effectively than that derived from DNABERT2.

In contrast, max pooling was associated with lower and more variable accuracies across cancer types, particularly for MLP and Naive Bayes models, indicating that retaining only the strongest token activations may discard relevant contextual information distributed throughout the sequence.

### 3.3 Model Assessment on Held-Out Datasets

The mean classification accuracy across cancer types was computed for each feature representation by averaging the accuracies of the seven models on the held-out test sets (Table 2 and Figure S6). Overall, features derived from gLM achieved the lowest average performance across datasets. NT and DNABERT exhibited greater variability among models, as indicated by the higher standard deviation values displayed above the bars. The k-mer and Word2Vec representations showed similar mean accuracies across datasets, suggesting stable overall performance. These results highlight the impact of representation strategies on model generalization and the importance of feature type selection in cancer-specific classification tasks.

**Table 2:** Mean (*μ*) and standard deviation (*σ*) of classification accuracy across the seven models for each feature representation and cancer dataset. Orange and blue values indicate the highest mean accuracy and lowest standard deviation, respectively, for each dataset.

| Datasets | DNABERT2 |  | KMER |  | NT |  | W2V |  |
| --- | --- | --- | --- | --- | --- | --- | --- | --- |
| | $\mu$ | $\sigma$ | $\mu$ | $\sigma$ | $\mu$ | $\sigma$ | $\mu$ | $\sigma$ |
| Anal Canal | 0.904 | 0.037 | **0.934** | **0.014** | **0.934** | 0.027 | 0.933 | 0.033 |
| Bile Duct | 0.929 | 0.016 | 0.942 | 0.016 | 0.887 | 0.027 | **0.952** | **0.011** |
| Bladder | 0.938 | 0.014 | 0.949 | **0.010** | 0.919 | 0.037 | **0.952** | 0.017 |
| Breast | 0.925 | 0.017 | 0.939 | **0.012** | 0.884 | 0.046 | **0.943** | 0.013 |
| Colon | 0.927 | 0.022 | 0.947 | 0.007 | 0.886 | 0.037 | **0.949** | **0.006** |
| Gastric | 0.892 | 0.054 | 0.936 | 0.020 | 0.899 | 0.048 | **0.938** | **0.017** |
| Kidney | 0.927 | 0.029 | 0.948 | 0.016 | 0.928 | 0.030 | **0.957** | **0.013** |
| Leukemia | 0.921 | 0.033 | 0.954 | 0.012 | 0.889 | 0.029 | **0.956** | **0.010** |
| Liver | 0.890 | 0.020 | 0.939 | **0.010** | 0.907 | 0.028 | **0.944** | 0.016 |
| Lung | 0.924 | 0.022 | 0.943 | **0.007** | 0.885 | 0.026 | **0.944** | 0.010 |
| Ovary | 0.954 | 0.024 | 0.972 | 0.021 | 0.924 | 0.035 | **0.978** | **0.011** |
| Prostate | 0.927 | 0.021 | 0.939 | 0.011 | 0.871 | 0.048 | **0.944** | **0.010** |
| Skin | 0.932 | 0.026 | **0.936** | **0.019** | 0.862 | 0.036 | 0.932 | 0.022 |
| Thyroid | 0.930 | 0.046 | 0.935 | 0.035 | 0.930 | 0.037 | **0.953** | **0.029** |
| Tongue | 0.922 | 0.033 | **0.948** | **0.014** | 0.915 | 0.026 | 0.943 | 0.032 |

To further investigate the influence of feature representation on model performance, we generated boxplots for each classifier to illustrate the distribution of mean accuracies across all cancer datasets for the four feature types (Figure 6). Word2Vec achieved the highest median accuracies for most models, particularly for Random Forest, Logistic Regression, XGBoost, and KNN. K-mer features showed comparable performance, especially when combined with SVM and MLP, confirming the strong predictive capacity of traditional sequence-based features. In contrast, the transformer-derived embeddings resulted in lower median accuracies and greater variability, particularly for KNN, Naive Bayes, and XGBoost. Among the gLM-based representations, DNABERT2 achieved higher median accuracies than NT. SVM and Logistic Regression were less affected by the choice of feature representation, exhibiting similar median accuracies across feature types, except for NT.

**Figure 6:**
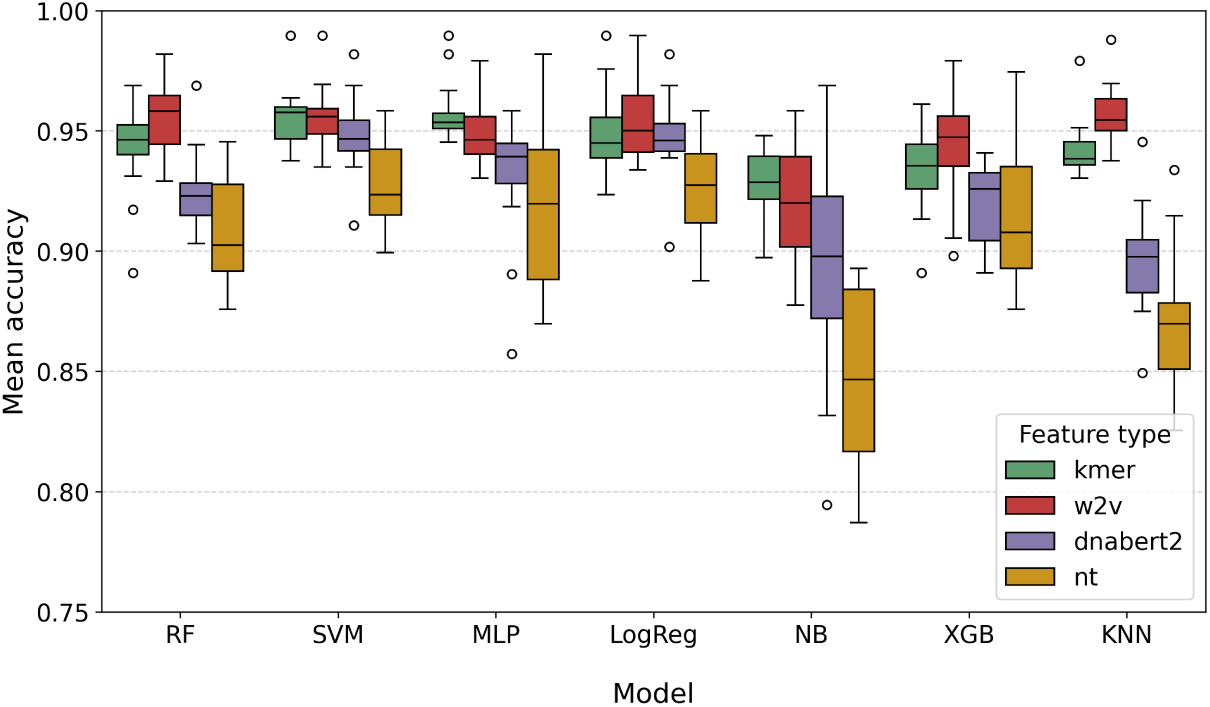
Distribution of mean classification accuracies across the 15 cancer-specific datasets for each machine learning model and feature representation (k-mer, Word2Vec, DNABERT2, and Nucleotide Transformer). Each boxplot summarizes the average performance obtained for a given model–feature combination.

The comparative analysis across all cancer types revealed consistent performance patterns among the evaluated feature representations and classification algorithms (Figure 7). The CoraL did not achieve the highest accuracy in any dataset, as the best predictive performance was consistently achieved by one of the evaluated feature–classifier combinations. Overall, models trained on k-mer and Word2Vec features achieved the highest accuracies across most datasets, reinforcing the strong predictive capability of these classical sequence representations. In particular, Word2Vec combined with SVM and KNN was the top-performing approach in 6 of the 15 datasets, while k-mer combined with SVM and MLP achieved the highest accuracy in 5 datasets each. Logistic Regression also demonstrated competitive performance, particularly when combined with Word2Vec and DNABERT2 embeddings, achieving the highest accuracy in 4 and 3 datasets, respectively. In contrast, Naive Bayes was outperformed by the remaining classifiers and did not achieve the best result in any dataset.

**Figure 7:**
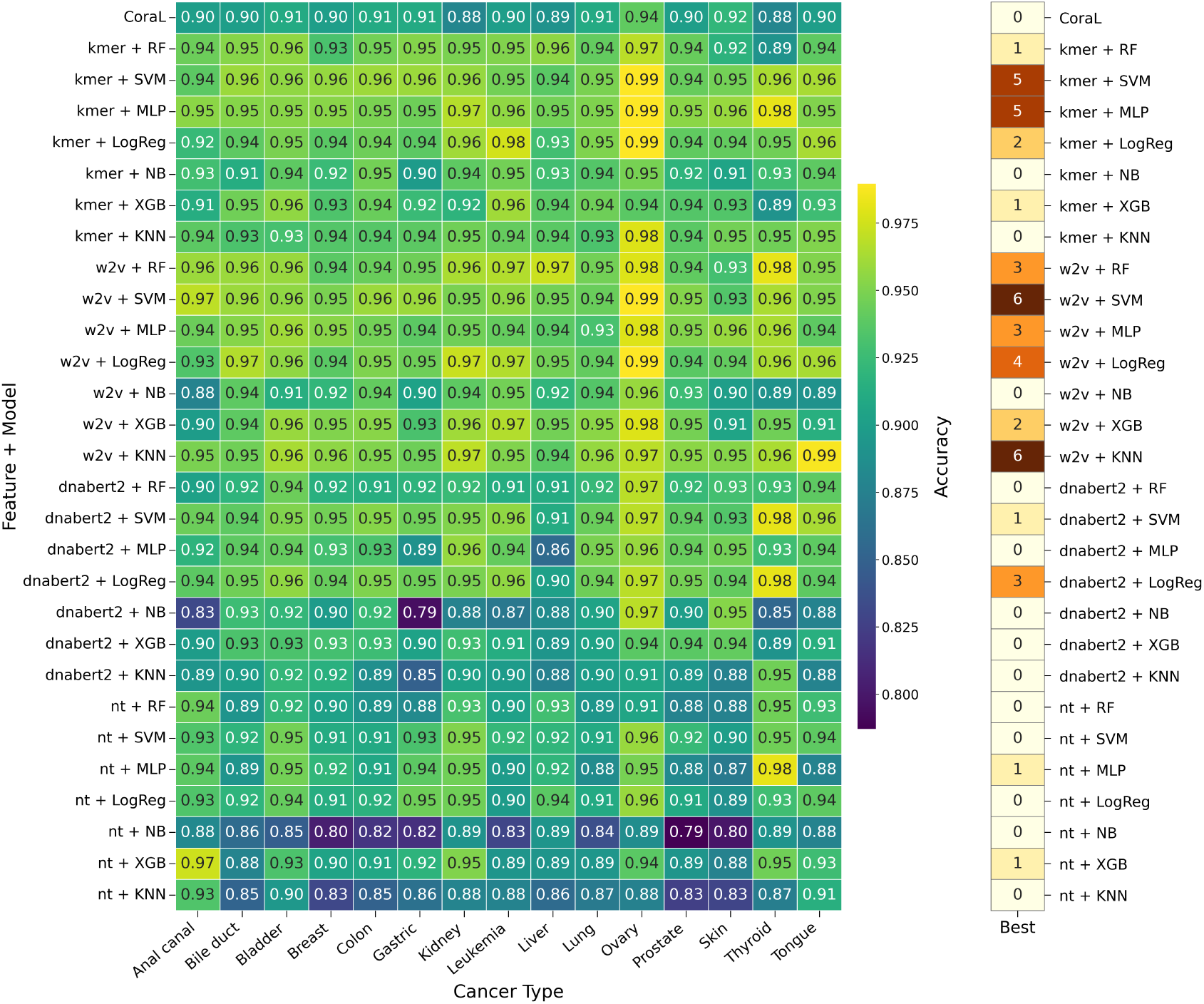
Heatmap of the classification accuracies across cancer types for all model–feature combinations. The ’Best’ column reports the number of times each model–pooling combination achieved the highest accuracy across the 15 datasets (ties counted for all winners).

Among the transformer-based representations, DNABERT2 outperformed NT and produced competitive results when combined with SVM and Logistic Regression. However, the gLM-based embeddings exhibited greater dependence on classifier selection than k-mer and Word2Vec representations. For example, in the gastric cancer dataset, DNABERT2 achieved accuracies ranging from 0.79 with Naive Bayes to 0.95 with Logistic Regression, whereas the corresponding Word2Vec-based models exhibited a difference of only 5%.

Finally, we assessed the performance of the models in terms of sensitivity and specificity, as accuracy alone can hide imbalances between identifying positive and negative cases (Figure 8). In general, most representations exhibit a reasonable trade-off between these two metrics, with some exceptions. For example, in the skin cancer dataset, we observed a consistent reduction in sensitivity, even for k-mer and Word2Vec representations (which show the best overall performance) when paired with Random Forest. This pattern suggests that certain datasets may pose intrinsic challenges that affect the models’ ability to detect positive cases. Overall, DNABERT2 combined with all classifiers, and particularly with Random Forest, showed a tendency toward reduced sensitivity. This indicates that, although DNABERT2 captures rich contextual information from sequences, its embeddings may be less effective at identifying positive instances in this specific problem setting. In contrast, NT representations exhibit a more balanced behavior regarding sensitivity and specificity, suggesting more stable performance across different decision boundaries.

**Figure 8:**
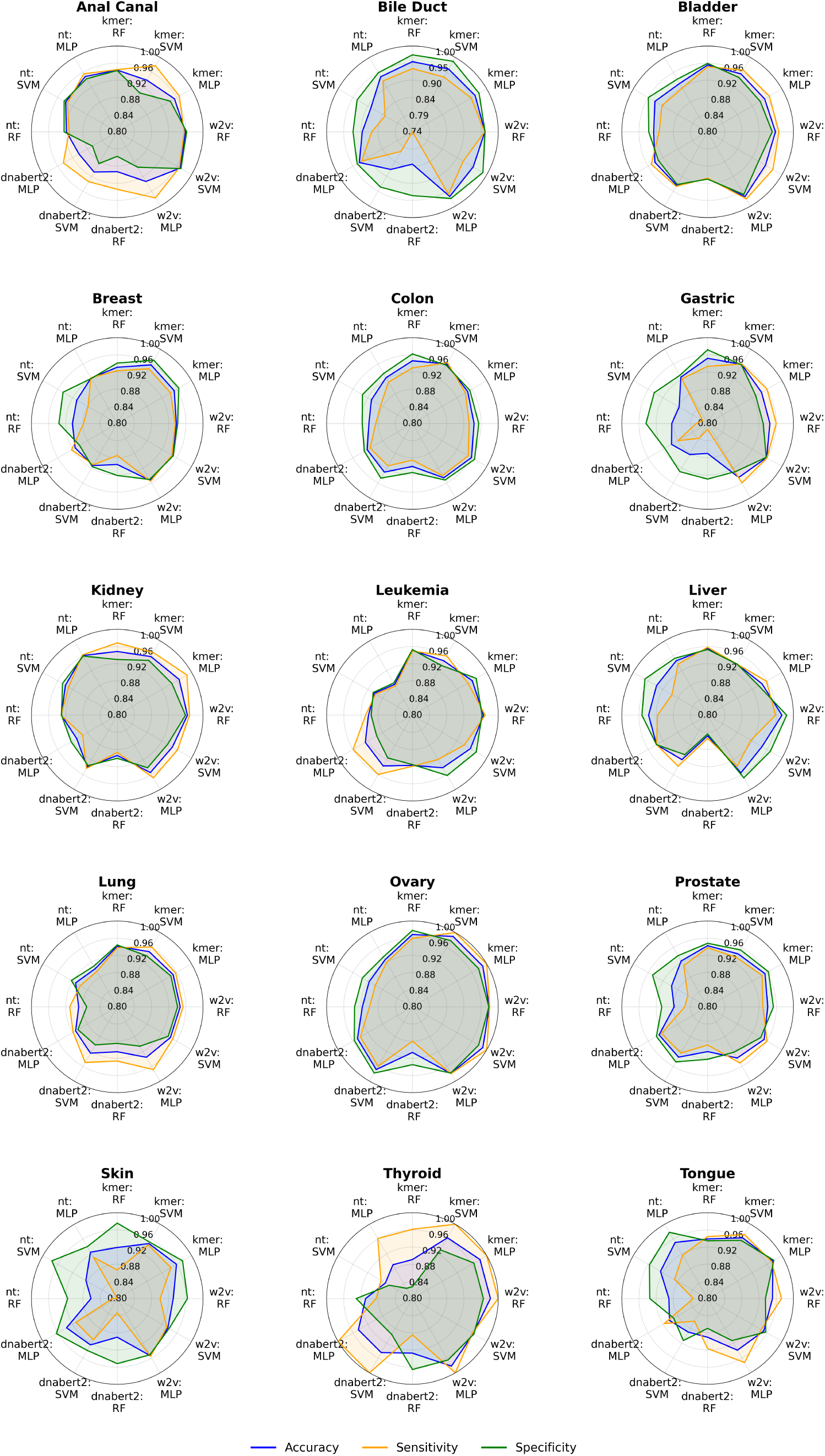
Evaluation of multiple feature–model combinations across cancer datasets using accuracy, sensitivity, and specificity.

## 4 Discussions

We presented a comprehensive evaluation of feature representation strategies for identifying cancer-associated sORFs in non-coding RNAs. Through an extensive benchmarking of seven supervised machine learning classifiers and three feature representation approaches across 15 cancer types, we identified the combinations that achieved the highest predictive performance for cancer-associated sORF prediction, an emerging research area in cancer genomics.

Our comparative analysis revealed that classical representations (k-mer frequency profiles and Word2Vec embeddings) consistently achieved superior accuracy compared to more complex gLM-based features across all datasets. These findings indicate that local sequence patterns capture discriminative patterns for the identification of cancer-associated sORF and that pre-trained genomic embeddings may require task-specific optimization. Furthermore, direct comparison with CoraL, the first published tool for cancer sORF prediction in non-coding RNAs, demonstrated superior performance of our framework across all tested datasets, emphasizing the importance of selecting feature representations and learning algorithms that are well aligned with the biological characteristics of the prediction task.

The datasets used in the experimental analysis were derived from those provided by CoraL, originally extracted from the SPENCER database, which compiles experimentally supported small peptides encoded by ncRNAs in cancer. A key limitation relates to the generation of negative data, which were obtained by shuffling positive sequences in the datasets introduced by CoraL, resulting in synthetic examples that may not fully represent the biological variability of truly non-functional sequences. As future work, we aim to explore more realistic strategies for defining negative data, as this remains an open problem that could substantially improve model robustness for real-world applications. Additionally, we intend to apply fine-tuning to the gLMs used in this study. Although these models already provide rich contextual representations, adapting them to task-specific data could further enhance their discriminative ability. Nevertheless, identifying the most suitable fine-tuning strategy is essential, particularly given that some datasets contain relatively few training samples, increasing the risk of overfitting and reducing the potential benefits of model adaptation.

Finally, this study shows that this pipeline could be applied to uncover biologically meaningful patterns in cancer-associated sORFs. For instance, it may help identify potential small peptides that could be expressed across multiple cancer types or those associated with clinical outcomes such as metastasis, disease progression, or response to therapy. Such analyses could support biomarker discovery, prioritize sORFs for experimental validation, and guide the identification of potential therapeutic targets. Beyond cancer, the approach could also be adapted to study peptides encoded by non-coding RNAs in other diseases, facilitating the exploration of their regulatory or signaling roles. Overall, these applications highlight the potential of integrative computational pipelines to connect sequence-level analyses with functional and translational biology.

## Acknowledgments

The authors acknowledge the Advanced Research Computing (ARC) team at the Rosalind Franklin Institute for the computational infrastructure and support. The Baskerville Tier 2 HPC service (https://www.baskerville.ac.uk/), which was funded by the EPSRC and UKRI: World Class Labs scheme (EP/T022221/1), the Digital Research Infrastructure programme (EP/W032244/1), and operated by ARC (University of Birmingham). The support of Health Data Research UK (HDR UK), funded by UK Research and Innovation, for its Black Internship Programme.

## References

[1] Kinga Nemeth, Recep Bayraktar, Manuela Ferracin, and George A. Calin. Non-coding rnas in disease: from mechanisms to therapeutics. Nature Reviews Genetics, 25(3):211–232, Mar 2024. ISSN 1471-0064. doi: 10.1038/s41576-023-00662-1. URL 10.1038/s41576-023-00662-1.

[2] Vinicius Maracaja-Coutinho, Alexandre Rossi Paschoal, José Carlos Caris-Maldonado, Pedro Vinícius Borges, Almir José Ferreira, and Alan Mitchell Durham. *Noncoding RNAs Databases: Current Status and Trends*, pages 251–285. Springer New York, New York, NY, 2019. ISBN 978-1-4939-8982-9. doi: 10.1007/978-1-4939-8982-9 10. URL 10.1007/978-1-4939-8982-9_10.

[3] Alexandre Rossi Paschoal, Vinicius Maracaja-Coutinho, João Carlos Setubal, Zilá Luz Paulino Simões, Sergio Verjovski-Almeida, and Alan Mitchell Durham. Non-coding transcription characterization and annotation. RNA Biology, 9(3):274–282, 2012. doi: 10.4161/rna.19352. PMID: 22336709.

[4] Sean R. Eddy. Non–coding rna genes and the modern rna world. Nature Reviews Genetics, 2(12): 919–929, Dec 2001. ISSN 1471-0064. doi: 10.1038/35103511. URL 10.1038/35103511.

[5] Nicholas T. Ingolia, Sina Ghaemmaghami, John R. S. Newman, and Jonathan S. Weissman. Genome-wide analysis in vivo of translation with nucleotide resolution using ribosome profiling. Science, 324(5924):218–223, 2009. doi: 10.1126/science.1168978. URL https://www.science.org/ doi/abs/10.1126/science.1168978.

[6] Ewen Callaway. ’dark proteins’ hiding in our cells could hold clues to cancer and other diseases. Nature, 637(8048):1038–1040, January 2025.

[7] Juan-Pablo Couso and Pedro Patraquim. Classification and function of small open reading frames. Nature Reviews Molecular Cell Biology, 18(9):575–589, Sep 2017. ISSN 1471-0080. doi: 10.1038/nrm.2017.58. URL 10.1038/nrm.2017.58.

[8] Jonathan M Mudge, Jorge Ruiz-Orera, John R Prensner, Marie A Brunet, Ferriol Calvet, Irwin Jungreis, Jose Manuel Gonzalez, Michele Magrane, Thomas F Martinez, Jana Felicitas Schulz, et al. Standardized annotation of translated open reading frames. Nature biotechnology, 40(7): 994–999, 2022.

[9] Micha l I. Świrski, Jack A. S. Tierney, M. Mar Albà, Dmitry E. Andreev, Julie L. Aspden, John F. Atkins, Michal Bassani-Sternberg, Marla J. Berry, Stefano Biffo, Kathleen Boris-Lawrie, Mark Borodovsky, Ian Brierley, Matthew Brook, Marie A. Brunet, Janusz M. Bujnicki, Neva Caliskan, Lorenzo Calviello, Anne-Ruxandra Carvunis, Jamie H. D. Cate, Can Cenik, Kung Yao Chang, Yi-wen Chen, Sonia Chothani, Jyoti S. Choudhary, Patricia L. Clark, Jim Clauwaert, Lynn Cooley, Erik Dassi, Kellie Dean, Jean-Jacques Diaz, Christoph Dieterich, Rivka Dikstein, Jonathan D. Dinman, Sergey E. Dmitriev, Olga A. Dontsova, Christine M. Dunham, Sandeep M. Eswarappa, Philip J. Farabaugh, Pouya Faridi, Ivo Fierro-Monti, Andrew E. Firth, David Gatfield, Fátima Gebauer, Mikhail S. Gelfand, Nicola K. Gray, Rachel Green, Chris H. Hill, Ya-Ming Hou, Norbert Hübner, Zoya Ignatova, Pavel Ivanov, Shintaro Iwasaki, Rory Johnson, Ahmad Jomaa, Marko Jovanovic, Irwin Jungreis, Manolis Kellis, Jeffrey S. Kieft, Alex V. Kochetov, Eugene V. Koonin, Andrei A. Korostelev, Joanna Kufel, Ivan V. Kulakovskiy, Leo Kurian, Denis L. J. Lafontaine, Ola Larsson, Gary Loughran, Julius Lukeš, Marco Mariotti, Elena S. Martens-Uzunova, Thomas F. Martinez, Akinobu Matsumoto, Joel McManus, Jan Medenbach, Sergey V. Melnikov, Gerben Menschaert, Catharina Merchante, Martin Mikl, W. Allen Miller, Oliver Mühlemann, Olivier Namy, Danny D. Nedialkova, Jozef Nosek, Sandra Orchard, Petar Ozretić, Mihaela Pertea, Dmitri D. Pervouchine, Lúısa Romão, David Ron, Xavier Roucou, Maria P. Rubtsova, Jorge Ruiz-Orera, Alan Saghatelian, Steven L. Salzberg, Lucia A. Seale, Cathal Seoighe, Petr V. Sergiev, Premal Shah, Nikolay Shirokikh, Sarah A. Slavoff, Nahum Sonenberg, Timothy J. Stasevich, Roman J. Szczesny, Tiina Tamm, Marek Tchórzewski, Ivan Topisirovic, Michel L. Tremblay, Tamir Tuller, Igor Ulitsky, Leoš Shivaya Valášek, Petra Van Damme, Gabriella Viero, Juan Antonio Vizcaino, Christine Vogel, Edward W. J. Wallace, Jonathan S. Weissman, Eric Westhof, Nicola Whiffin, Daniel N. Wilson, Zhi Xie, Jonathan W. Yewdell, Martina M. Yordanova, Chien-Hung Yu, Vyacheslav Yurchenko, Bojan Zagrovic, Maria Inês Almeida, Nese Atabey, Nikolaos Balatsos, Pavel Baranov, Anca Simona Bojan, Theodora Choli-Papadopoulou, Pierre Close, Victoria Cowling, Alexandre David, Aleksandar Ef-timov, Mark Helm, Cristina-Adela Iuga, Dana Jurkovičová, Arvydas Kanopka, Denis Lafontaine, Elena Martens-Uzunova, Lucia Messingerová, Henrik Nielsen, Mehmet Ozturk, Vicent Pelechano, Marianna Penzo, Paulina Podszywalow-Bartnicka, Lejla Pojskić, John Le Quesne, Barak Rotblat, Ariel Stanhill, Georg Stoecklin, Marek Tchorzewski, Vladimir Trajković, Leos Shivaya Valasek, Eivind Valen, Sinisa Volarevic, Ljubica Vucicevic, Kathleen Watt, Anne Willis, Masa Zdrale-vic, Taja Železnik Ramuta, Pavel V. Baranov, and TRANSLACORE. Translon: a single term for translated regions. Nature Methods, 22(10):2002–2006, Oct 2025. ISSN 1548-7105. doi: 10.1038/s41592-025-02810-3. URL 10.1038/s41592-025-02810-3.

[10] SURESH C. GOEL. Transcription unit. Nature, 245(5425):397–397, Oct 1973. ISSN 1476-4687. doi: 10.1038/245397c0. URL 10.1038/245397c0.

[11] Diego Guerra-Almeida, Diogo Antonio Tschoeke, and Rodrigo Nunes-da Fonseca. Understanding small orf diversity through a comprehensive transcription feature classification. DNA Research, 28 (5):dsab007, 2021.

[12] D Guerra-Almeida and R Nunes-da Fonseca. Small open reading frames: how important are they for molecular evolution? front. genet. 11, 574737, 2020.

[13] Julie L Aspden, Ying Chen Eyre-Walker, Rose J Phillips, Unum Amin, Muhammad Ali S Mumtaz, Michele Brocard, and Juan-Pablo Couso. Extensive translation of small open reading frames revealed by poly-ribo-seq. elife, 3:e03528, 2014.

[14] Yaguang Zhang. Lncrna-encoded peptides in cancer. Journal of Hematology & Oncology, 17(1): 66, 2024.

[15] Jizhong Wang, Song Zhu, Nan Meng, Yutian He, Ruixun Lu, and Guang-Rong Yan. ncrna-encoded peptides or proteins and cancer. Molecular Therapy, 27(10):1718–1725, 2019.

[16] Gregory Tong and Thomas F. Martinez. Ribosome profiling reveals hidden world of small proteins. Trends in Genetics, 41(2):101–103, 2025. ISSN 0168-9525. doi: 10.1016/j.tig.2024.12.010. URL https://www.sciencedirect.com/science/article/pii/S0168952524003184. Special issue: Microproteins.

[17] Joshua Beals, Haiyan Hu, and Xiaoman Li. A survey of experimental and computational identification of small proteins. Briefings in Bioinformatics, 25(4):bbae345, 2024.

[18] Mengmeng Zhu and Michael Gribskov. Mipepid: Micropeptide identification tool using machine learning. BMC bioinformatics, 20(1):559, 2019.

[19] Meng Zhang, Jian Zhao, Chen Li, Fang Ge, Jing Wu, Bin Jiang, Jiangning Song, and Xiaofeng Song. csorf-finder: an effective ensemble learning framework for accurate identification of multi-species coding short open reading frames. Briefings in Bioinformatics, 23(6), 2022.

[20] Zhao Peng, Jiaqiang Li, Xingpeng Jiang, and Cuihong Wan. socp: a framework predicting smorf coding potential based on tis and in-frame features and effectively applied in the human genome. Briefings in Bioinformatics, 25(3):bbae147, 2024.

[21] Zhongshen Li, Junru Jin, Wenjia He, Wentao Long, Haoqing Yu, Xin Gao, Kenta Nakai, Quan Zou, and Leyi Wei. Coral: interpretable contrastive meta-learning for the prediction of cancer-associated ncrna-encoded small peptides. Briefings in Bioinformatics, 24(6):bbad352, 2023.

[22] Yoon Kim. Convolutional neural networks for sentence classification. arXiv preprint arXiv:1408.5882, 2014.

[23] Edo Dotan, Gal Jaschek, Tal Pupko, and Yonatan Belinkov. Effect of tokenization on transformers for biological sequences. Bioinformatics, 40(4):btae196, 2024.

[24] Tin Kam Ho. Random decision forests. In *Proceedings of 3rd international conference on document analysis and recognition*, volume 1, pages 278–282. IEEE, 1995.

[25] Vladimir Vapnik. Support-vector networks. Machine learning, 20:273–297, 1995.

[26] David E Rumelhart, Geoffrey E Hinton, and Ronald J Williams. Learning representations by back-propagating errors. nature, 323(6088):533–536, 1986.

[27] Xiaotong Luo, Yuantai Huang, Huiqin Li, Yihai Luo, Zhixiang Zuo, Jian Ren, and Yubin Xie. Spencer: a comprehensive database for small peptides encoded by noncoding rnas in cancer patients. Nucleic acids research, 50(D1):D1373–D1381, 2022.

[28] Tomas Mikolov, Kai Chen, Greg Corrado, and Jeffrey Dean. Efficient estimation of word representations in vector space. arXiv preprint arXiv:1301.3781, 2013.

[29] Zhihan Zhou, Yanrong Ji, Weijian Li, Pratik Dutta, Ramana Davuluri, and Han Liu. Dnabert-2: Efficient foundation model and benchmark for multi-species genome. arXiv preprint arXiv:2306.15006, 2023.

[30] Hugo Dalla-Torre, Liam Gonzalez, Javier Mendoza-Revilla, Nicolas Lopez Carranza, Adam Henryk Grzywaczewski, Francesco Oteri, Christian Dallago, Evan Trop, Bernardo P de Almeida, Hassan Sirelkhatim, et al. Nucleotide transformer: building and evaluating robust foundation models for human genomics. Nature Methods, 22(2):287–297, 2025.

