## Supplementary Material for "Systematic Evaluation of Feature Representations for Cancer-Associated sORF Prediction in Non-coding RNA"

### Supplementary Figures

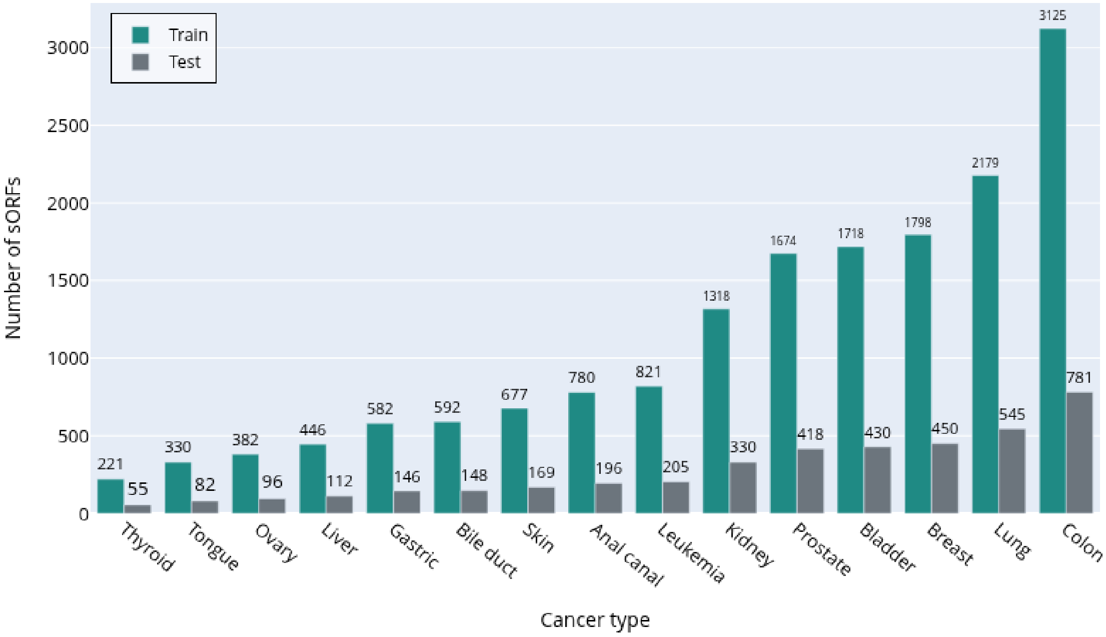

**Figure S1.** CoraL datasets statistics.

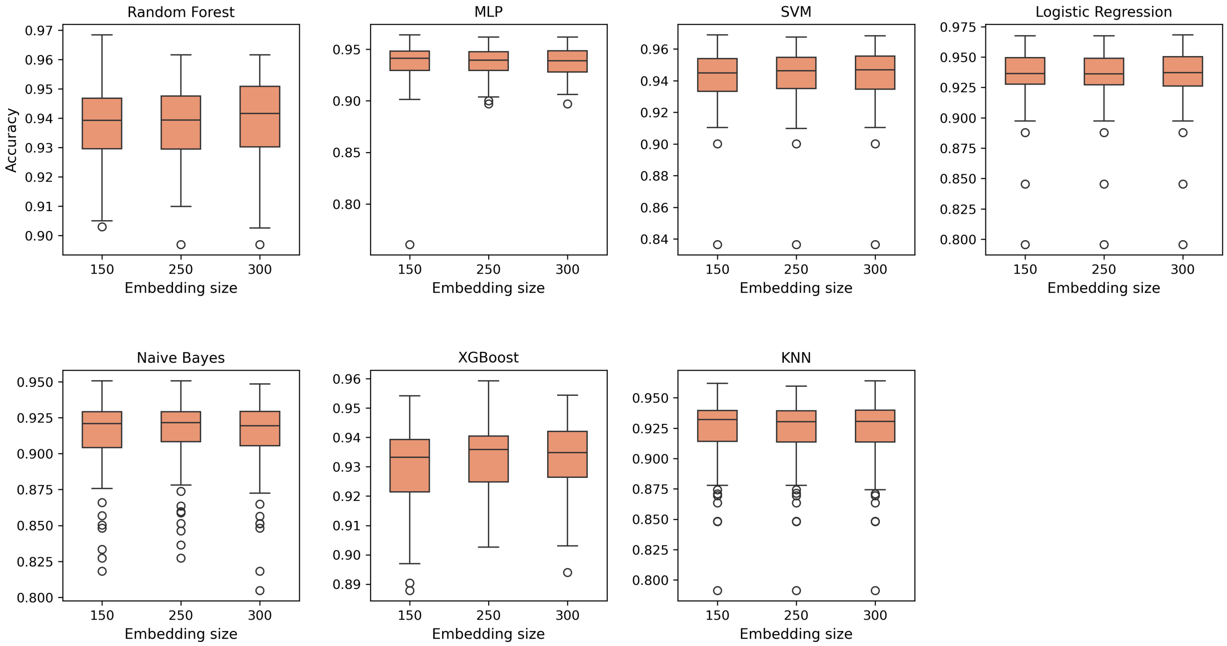

**Figure S2.** Hyperparameter performance analysis. Word2Vec embedding vector sizes under varying hyperparameter settings.

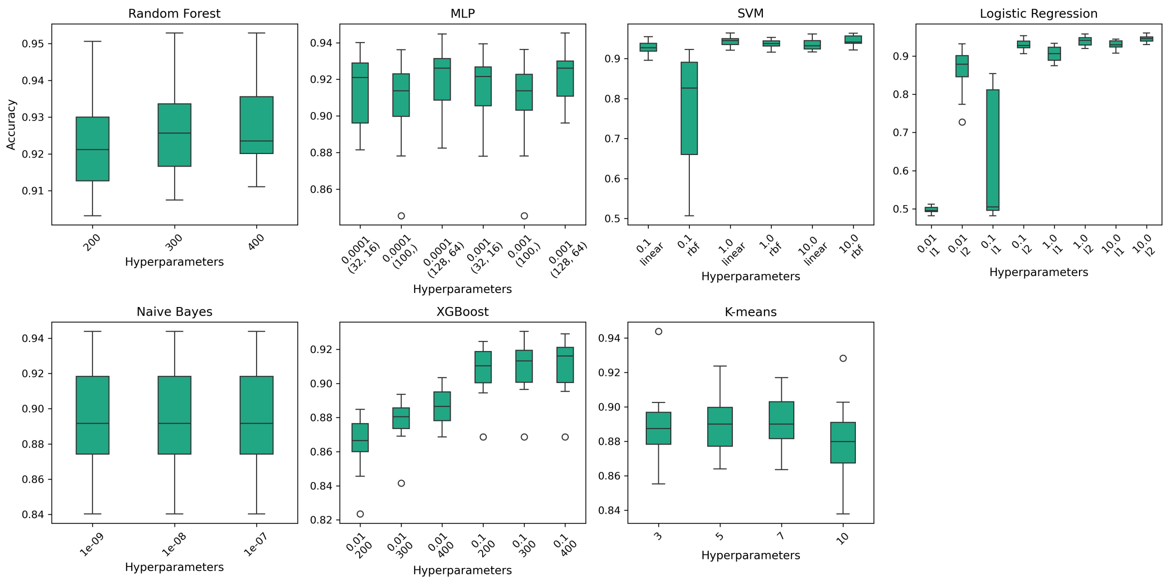

**(a)** Mean pooling

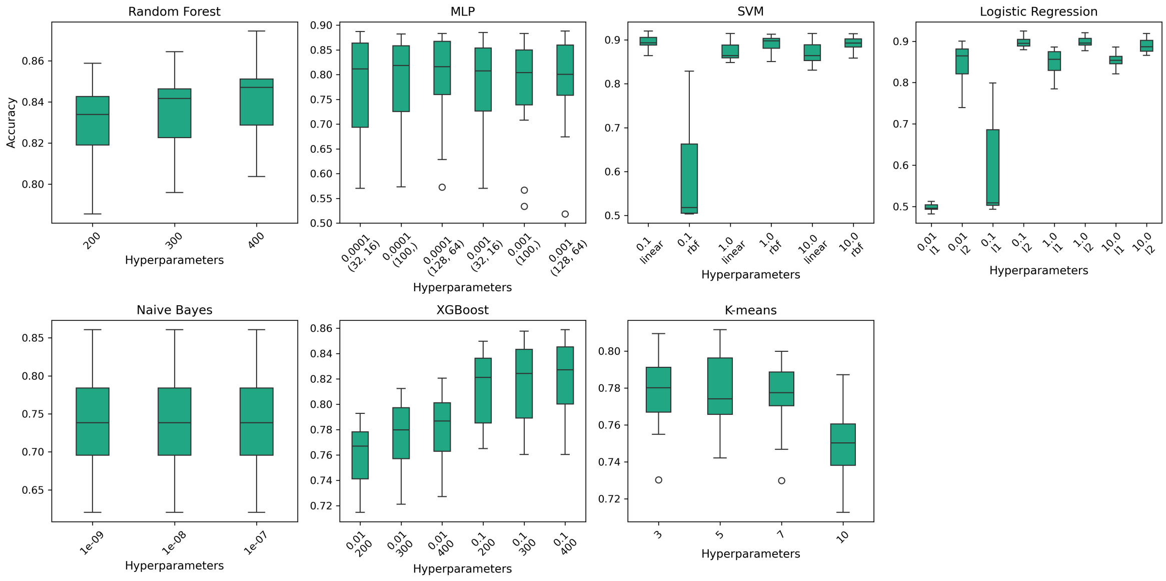

**(b)** Max pooling

**Figure S3.** Hyperparameter performance analysis. Boxplots show the mean cross-validation accuracies for each classifier using DNABERT2 with (a) mean and (b) max pooling under varying hyperparameter settings.

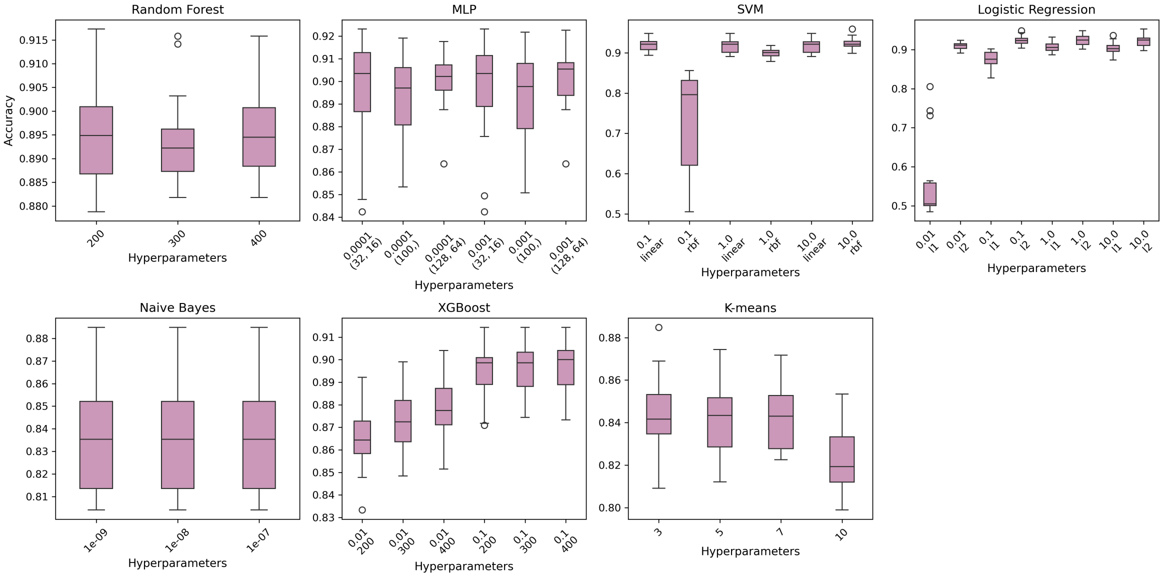

**(a)** Mean pooling

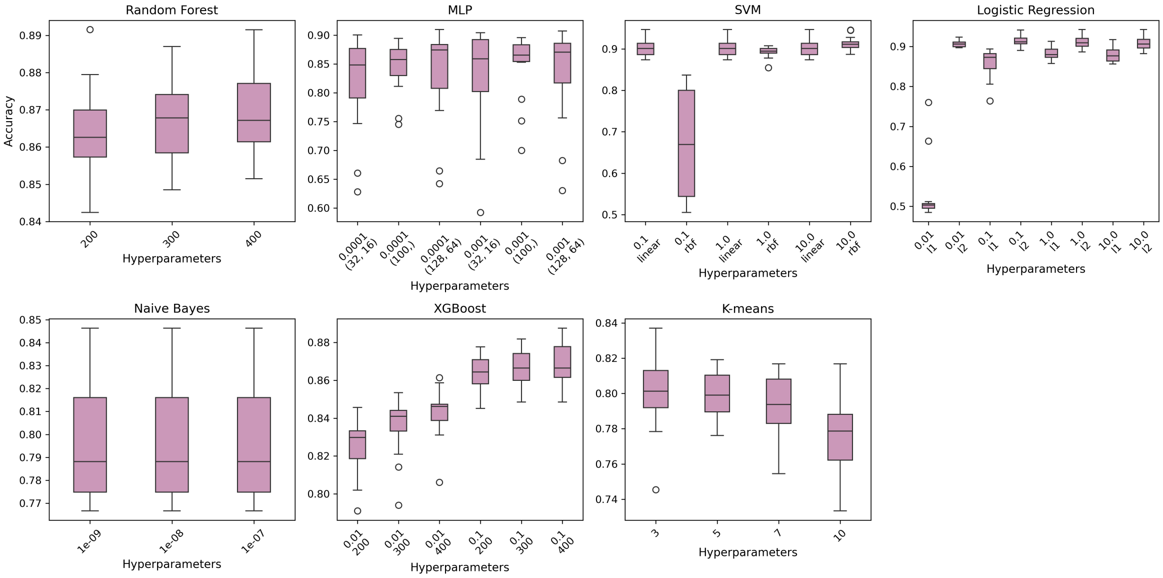

**(b)** Max pooling

**Figure S4.** Hyperparameter performance analysis. Boxplots show the mean cross-validation accuracies for each classifier using Nucleotide Transformer with (a) mean and (b) max pooling under varying hyperparameter settings.

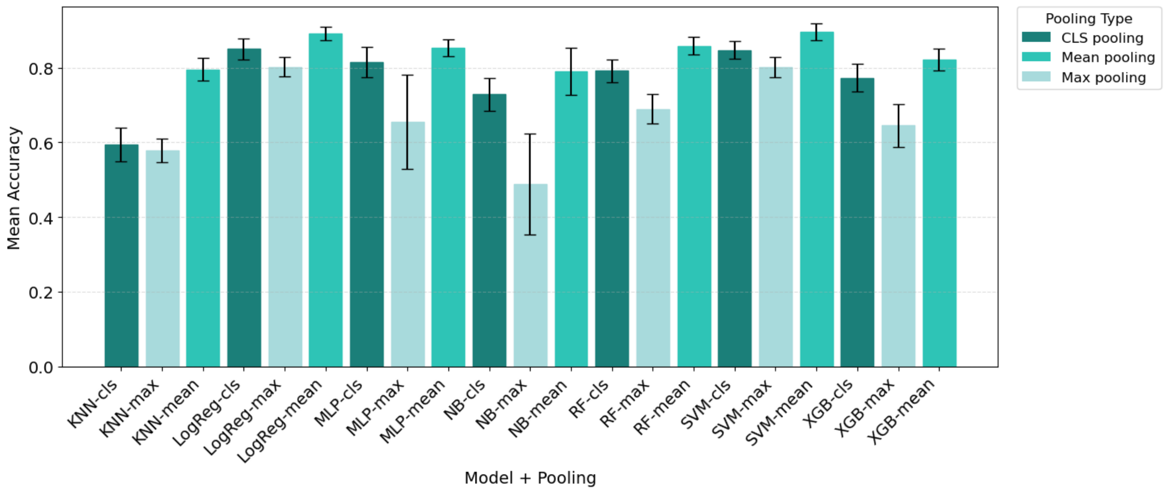

**(a)** DNABERT2

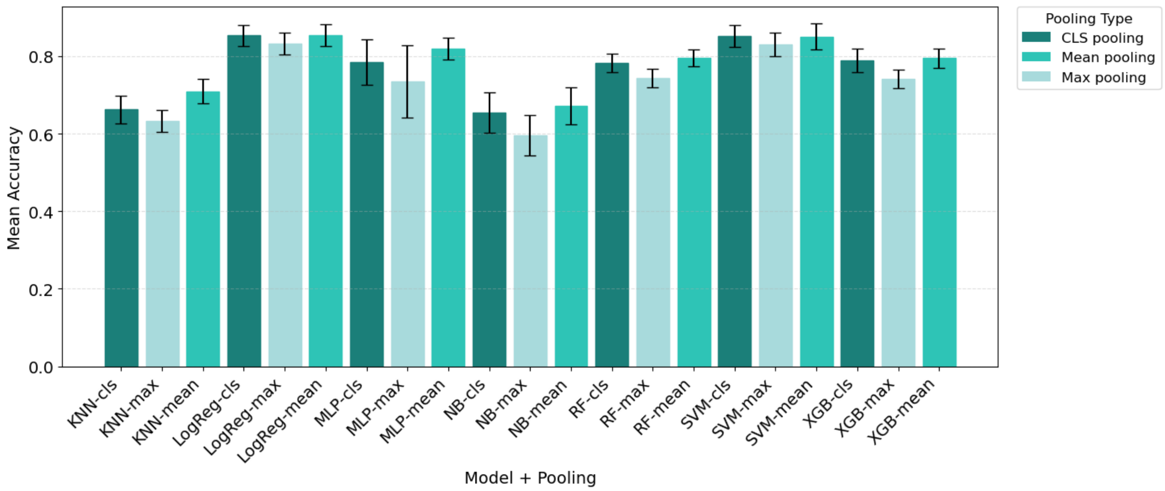

**(b)** Nucleotide Transformer

**Figure S5.** Mean classification accuracy across the 15 cancer datasets for seven machine learning models combined with three embedding pooling strategies. Bars represent the average accuracy obtained across all cancer datasets, and error bars indicate the standard deviation. Figures (a) and (b) show the results obtained with DNABERT2 and NT, respectively.

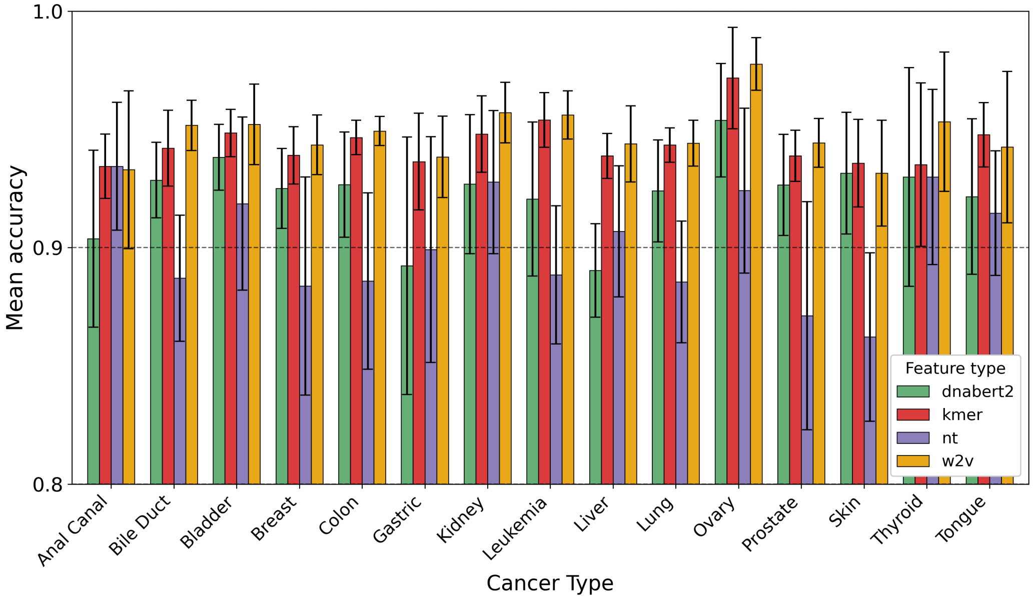

**Figure S6.** Mean classification accuracy (± standard deviation) across seven models for each feature representation, evaluated on held-out test datasets from 15 cancer types.
